# Chronic Intermittent Hypobaric Hypoxia Prevents Contrast-Induced Acute Kidney Injury By Modulating The HIF-1α Signaling Pathway

**DOI:** 10.1101/2022.02.07.479411

**Authors:** Kai-min Yin, Yan-hui Ni, Guang-yun Cao, Jia-yuan Zhang, Bao-jun Yi, Zi-hao Pang, Hui-jie Ma, Li Zhang

**Author notes:** Corresponding author: Li Zhang, Department of Cardiology, Hebei General Hospital, 348 West He Ping Road, Shijiazhuang, Hebei 050000, China, E-mail addresses (L. Zhang).

## Abstract

The aim of this study was to explore the role of CIHH in preventing contrast-induced acute kidney injury (CI-AKI) in rats and its mechanism. Rats mean arterial pressure, heart rate, serum creatinine and blood urea nitrogen levels were measured. The kidney tissue pathological changes, superoxide dismutase (SOD) activity, malondialdehyde (MDA) levels, hypoxia inducible factor-1α, Bcl-2/adenovirus E1B-19kDa-interacting protein3 (BNIP3) , cysteiny aspartate specific protease3(caspase3) and poly(ADP-ribose) polymerase (PARP) expression levels were testing. The results showed that CIHH prevented CI-AKI group mean arterial pressure, heart rate, serum creatinine and blood urea nitrogen levels were reduced, kidney tissue SOD activity was increased, MDA levels was reduced, HIF-1α,BNIP3,caspase3 and PARP levels were increased than the CI-AKI group. This study indicates that CIHH pretreatment may have a protective effect on contrast-induced early kidney injury by activating the HIF-1α/BNIP3 signaling pathway to regulate mitochondrial autophagy and enhance cellular anti-apoptotic and renal antioxidant capacity, for the first time.

## 1. Introduction

Although the incidence of CI-AKI has been reduced by the application of low osmotic / isotonic nonionic contrast agents, the incidence rate of CI-AKI is increasing due to the increase in high-risk groups such as elderly patients, diabetes mellitus and chronic kidney disease. It has been reported that CI-AKI has become the third leading cause of medically induced acute kidney injury[1].Once contrast renal damage occurs, there is still no effective treatment. Therefore, it is particularly important to prevent the occurrence of CI-AKI. The pathogenesis of CI-AKI is still unclear, but studies have shown that there are multiple potential pathways of interaction in the CI-AKI happen and development, including ischemia, hypoxia and oxidative stress due to altered renal hemodynamics caused by vasoactive substances and the high viscosity of the contrast agent itself, and direct toxic effects of the contrast agent itself on renal tubular epithelial cells and endothelial cells[2].

SOD has the function of scavenging ROS, after receiving contrast agent, the kidney undergoes oxidative stress, the activity of SOD in serum and kidney tissue decreases, the level of ROS and MDA increases, and ROS can directly cause tubular cell damage[3].Under hypoxia, the body activates hypoxia inducible factor (HIF) to regulate the expression of various genes to adapt to the hypoxic environment, such as genes involved in cell survival, angiogenesis, glycolysis, and invasion/metastasis[4]. HIF consists of three main α subunits (HIF-1α, HIF-2α, HIF-3α) and one β subunit (HIF-1β), HIF-1α and HIF-2α will becoming the main regulatory factors produced under hypoxic conditions[5]. It was shown that the expression levels of HIF-1α and BNIP3 were increased in contrast-induced acute kidney injury, and further increase of both expression could enhance mitochondrial autophagy and attenuate apoptosis to improve contrast kidney injury[6–8]. Caspase-3 is the major terminal shear enzyme in apoptosis, and its main substrate is PARP. When the body was stimulated, caspase3 was activated into cleaved-caspase3, cleaved-caspase3 could activate PARP to induce apoptosis. Romano et al study shows, human embryonic kidney cells (HEK 293), porcine proximal renal tubular cells (LLC-PK1) and Madin-Darby renal epithelial cells (MDCK) were incubated with Iodine contrast agent injections, caspase3 and RARP levels were reduced, cell activity was reduced, and apoptosis was increased, suggesting that iodine contrast agents can activate the caspase3 pathway to cause apoptosis in kidney cells[9]. Studies mentioned above showed that modulation of HIF-1α signaling pathway and inhibition of caspase-3 activation could enhance mitochondrial autophagy and attenuate apoptosis to protect the kidney. Thus, the changes of HIF-1α and caspase-3 levels were used as the main observation indicators in this experiment.

Current studies have shown the existence of multiple benefits of CIHH to the organism. For example, CIHH ameliorates renal injury in rats with diabetic nephropathy by activating the HIF1-α signaling pathway[10]; intermittent hypobaric hypoxia enhances HIF stability by inhibiting prolyl hydroxylase (PHD) activity, which activates key adaptive genes to attenuate hypoxic injury in the body[11],CIHH also has the ability to improve hypertension[12, 13], improve left ventricular remodeling and myocardial fibrosis[14],anti-arrhythmia[15, 16], attenuate oxidative stress[17], alter cardiac energy metabolism[18], reduce the size of myocardial infarction[19], inhibit inflammatory response and apoptosis[20], improve insulin resistance[13], attenuate endoplasmic reticulum stress[21].

Therefore, we hypothesized that CIHH may protect against CI-AKI by modulating the HIF signaling pathway, inhibiting renal oxidative stress, and enhancing mitochondrial autophagy and cellular anti-apoptotic capacity. In this study, we used histological and molecular biological methods to assess the protective effects of CIHH on CI-AKI rats induced by indomethacin, Nω-nitro-L-arginine methyl ester and ioversol injection, and investigate the mechanism of action4.

## 2. Results

### 2.1 Effect of CIHH on Heart Rate and Mean Arterial Pressure

One week before injection, all rats heart rate and mean arterial pressure were not difference. However, 24 hours after contrast agent injection, the heart rate and mean arterial pressure were significantly higher in the CI-AKI and CIHH prevented CI-AKI groups compared with the CTL group, and were significantly lower in the CIHH prevented CI-AKI group compared with the CI-AKI group (P<0.05, Fig 1).

**Figure1:**
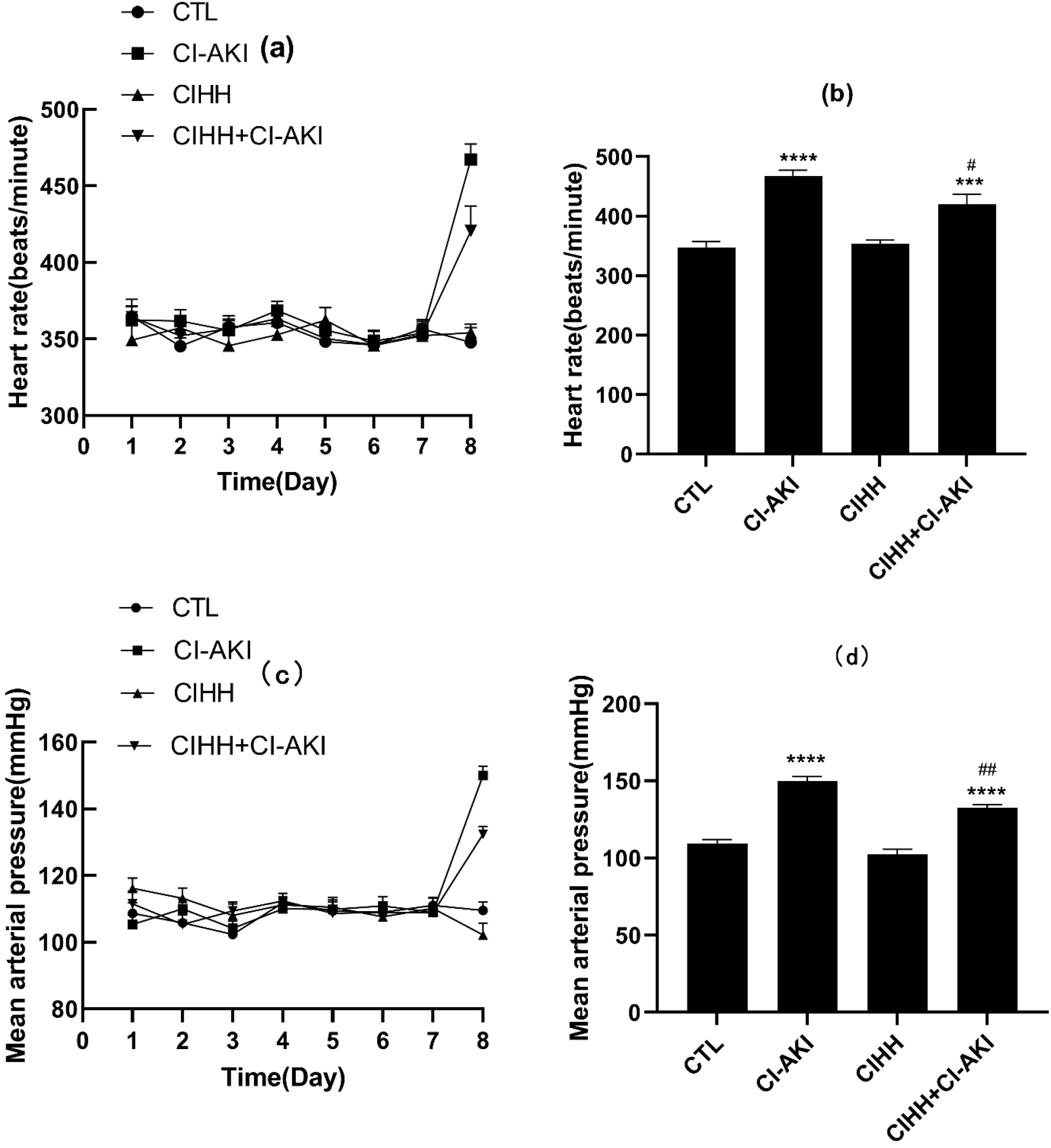
Effects of CIHH on heart rate and mean arterial pressure. The heart rate and mean arterial pressure dates were collected from seven days before injection to 24 hours after injection. The heart rate dates from one day to seven days before injection and 24 hours after injection(a), the heart rate dates at 24 hours after injection (b). The mean arterial pressure dates from one day to seven days before injection and 24 hours after injection (c), the mean arterial pressure dates at 24 hours after injection (d). CTL, normal-controlled group; CI-AKI, contrast induced acute kidney injury group; CIHH, chronic intermittent hypobaric hypoxia pretreatment group; CIHH+CI-AKI, contrast induced acute kidney injury group with chronic intermittent hypobaric hypoxia pretreatment. Data were presented as means ± SEM (n=6). ***P<0.001, ****p<0.0001versus.CTL; #p< 0.05, ##P< 0.01 versus.CI-AKI.

### 2.2 Effect of CIHH on renal function

Serum creatinine and blood urea nitrogen levels were significantly higher in the CI-AKI group and CIHH prevented CI-AKI group compared with the CTL group, but lower in the CIHH prevented CI-AKI group compared with the CI-AKI group (P<0.05, Fig 2).

**Figure 2:**
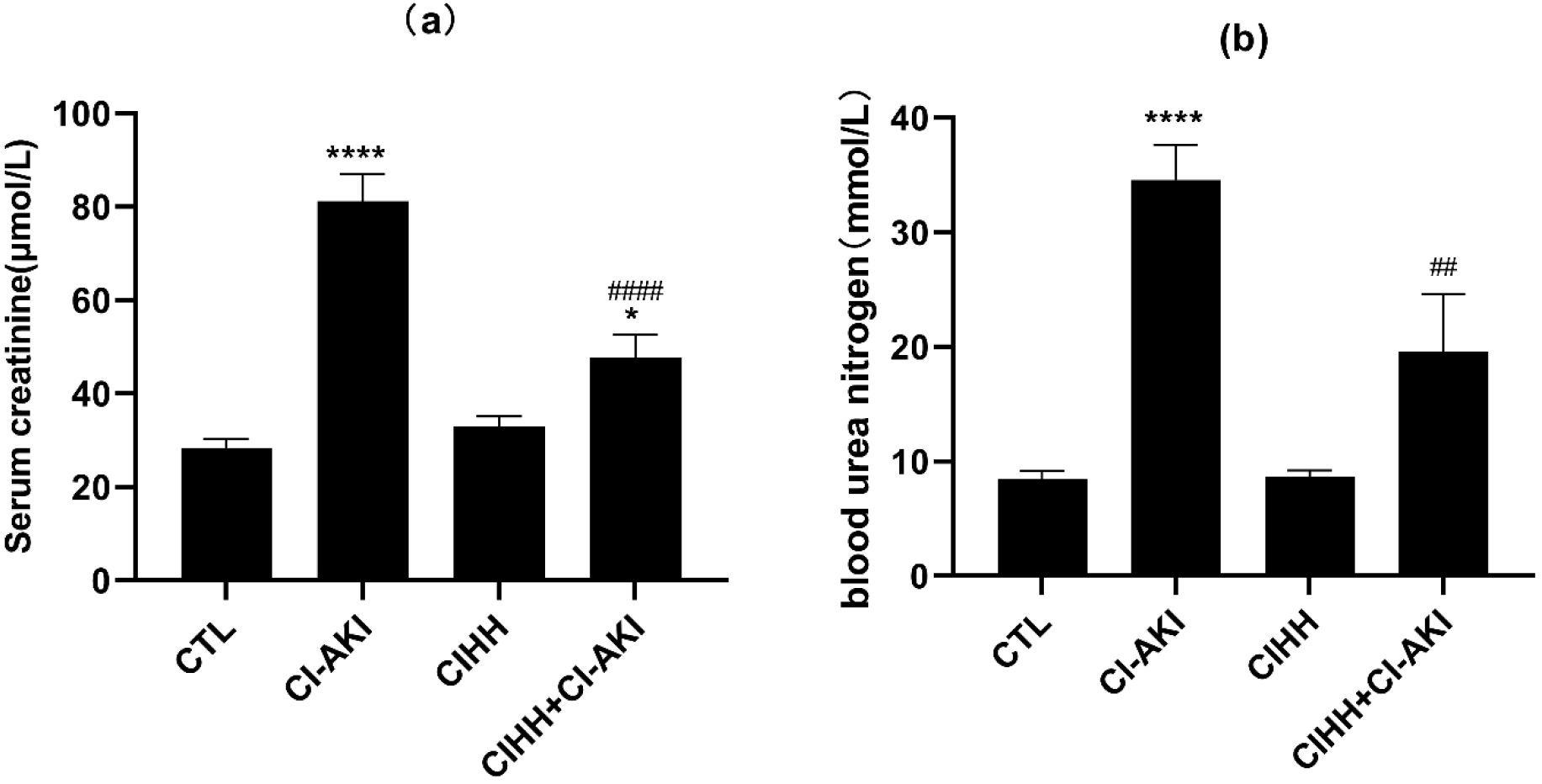
Effects of CIHH on renal function. Blood samples were collected through caudal vein before and 24 hours after contrast medium injection. Serum creatinine (a), blood nitrogen urea (b). CTL, normal-controlled group; CI-AKI, contrast induced acute kidney injury group; CIHH, chronic intermittent hypobaric hypoxia pretreatment group; CIHH+CI-AKI, contrast induced acute kidney injury group with chronic intermittent hypobaric hypoxia pretreatment. Data were presented as means ± SEM (n=6). *P<0.05, ****P<0.0001 versus CTL; ##p< 0.01, ####p< 0.0001versusCI-AKI.

### 2.3 Histological examination

HE staining showed that the water like degeneration of renal tubular epithelial cells in CI-AKI group was obvious, eosinophilic masses were seen in a large number of renal tubular lumens, some renal tubules were necrotic, and the lumen structure of renal tubules was not clear. There was a small amount of watery degeneration of renal tubular epithelial cells in CIHH prevented CI-AKI group, a small amount to medium amounts of eosinophilic masses in the lumen of renal tubules, and no tubular necrosis and changes in lumen structure were observed. PAS staining showed no obvious changes in glomerulus mesangial region, and Masson staining showed no obvious fibrosis changes in glomerulus and renal stroma in all groups.

**Fig3:**
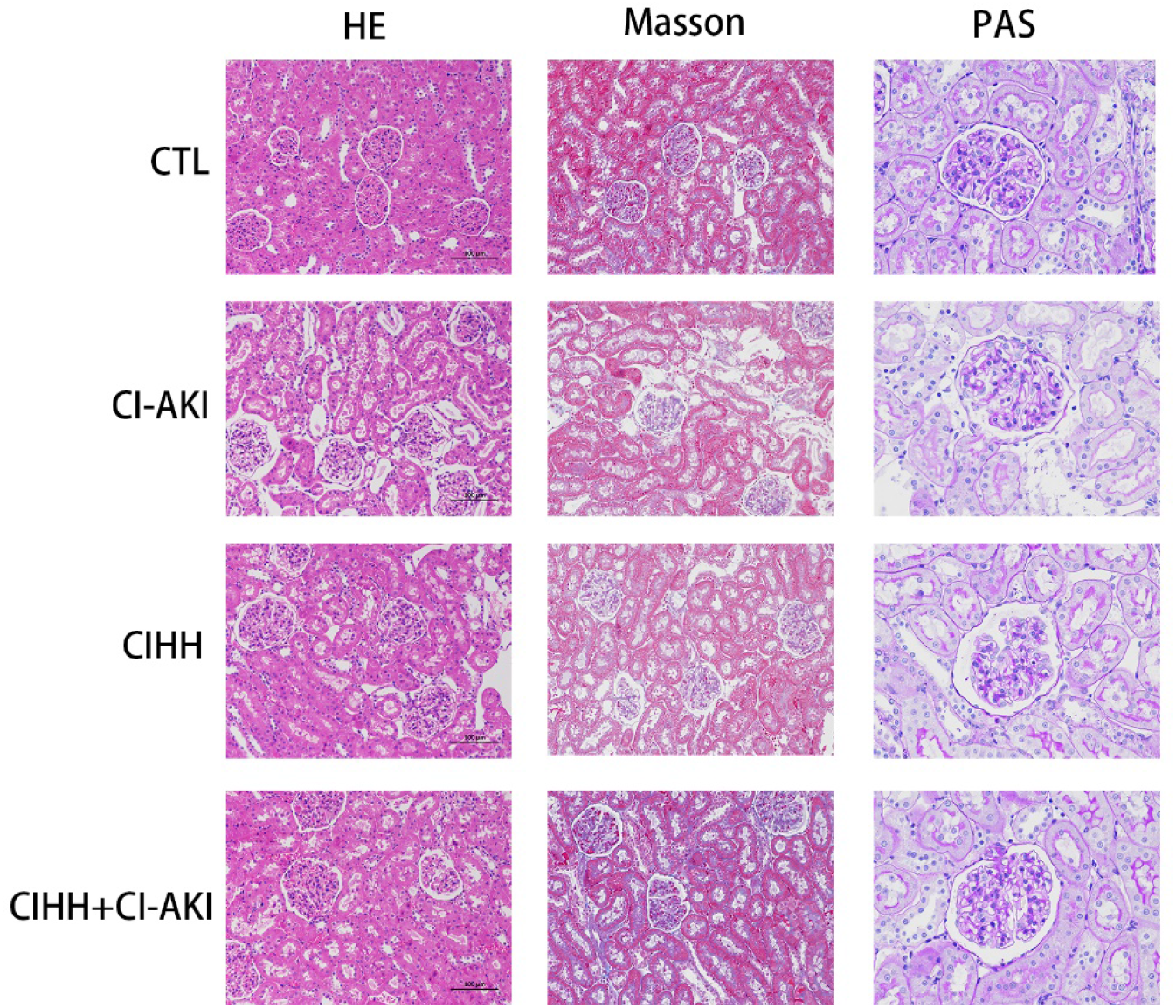
Effects of CIHH on renal histological examination. HE, hematoxylin eosin staining; PAS, periodic acid Schiff reaction; Masson, Masson trichrome. CTL, normal-controlled group; CI-AKI, contrast induced acute kidney injury group; CIHH, chronic intermittent hypobaric hypoxia pretreatment group; CIHH+CI-AKI, contrast induced acute kidney injury group with chronic intermittent hypobaric hypoxia pretreatment.

### 2.4 Antioxidant effects of CIHH

The activity of SOD was significantly decreased in CI-AKI group compared to CTL group, but was significantly increased in CIHH prevented CI-AKI group compared to CI-AKI group (P < 0.05, Fig. 4a). In addition, the levels of MDA were significantly higher in CI-AKI group compared to CTL group, but were significantly lower in CIHH prevented group compared to CI-AKI group (P < 0.05, Fig. 4b).

**Fig.4:**
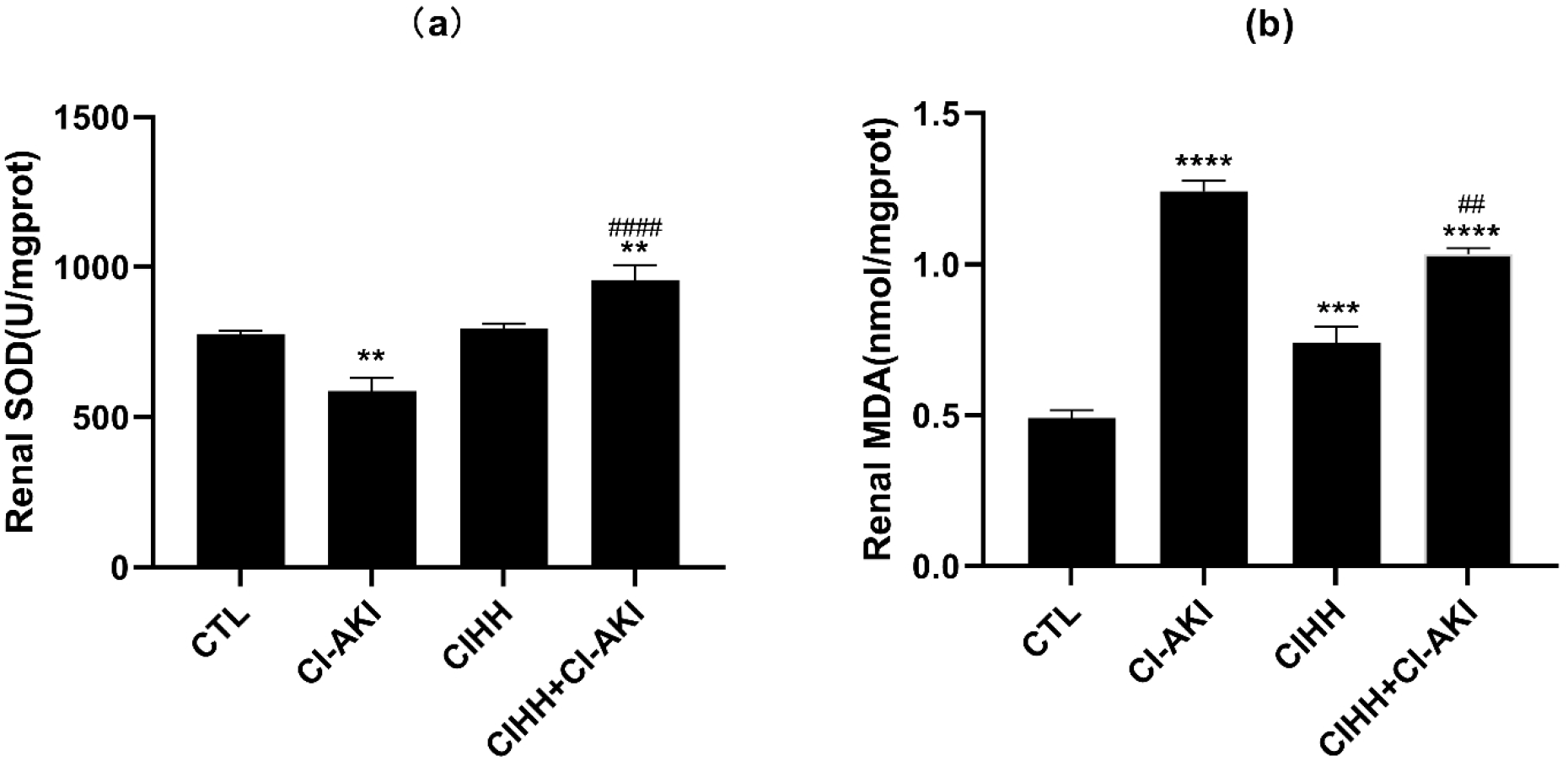
Effects of CIHH on renal oxidative stress. The levels of malondialdehyde (MDA) and superoxide dismutase (SOD) were measured to determine the alteration of oxidative stress. Renal MDA(a), Renal SOD(b). CTL, normal-controlled group; CI-AKI, contrast induced acute kidney injury group; CIHH, chronic intermittent hypobaric hypoxia pretreatment group; CIHH+CI-AKI, contrast induced acute kidney injury group with chronic intermittent hypobaric hypoxia pretreatment. Dates were presented as means ± SEM (n=6).**P<0.01, ***P<0.001,****P<0.0001versus.CTL, ##p< 0.01,####p< 0.0001versus.CI-AKI.

### 2.5 Western Blot

Our dates indicated that HIF-1α and BNIP3 protein expression levels were significantly increased in CI-AKI group compared to CTL group, and were more increased in CIHH prevented CI-AKI group compared to CI-AKI group. The results also showed that caspase3 and PARP levels were both down-regulated in CI-AKI group compared to CTL group, while increased inCIHH prevented CI-AKI group compared to CI-AKI group (P < 0.05, Fig. 5).

**Figure 5:**
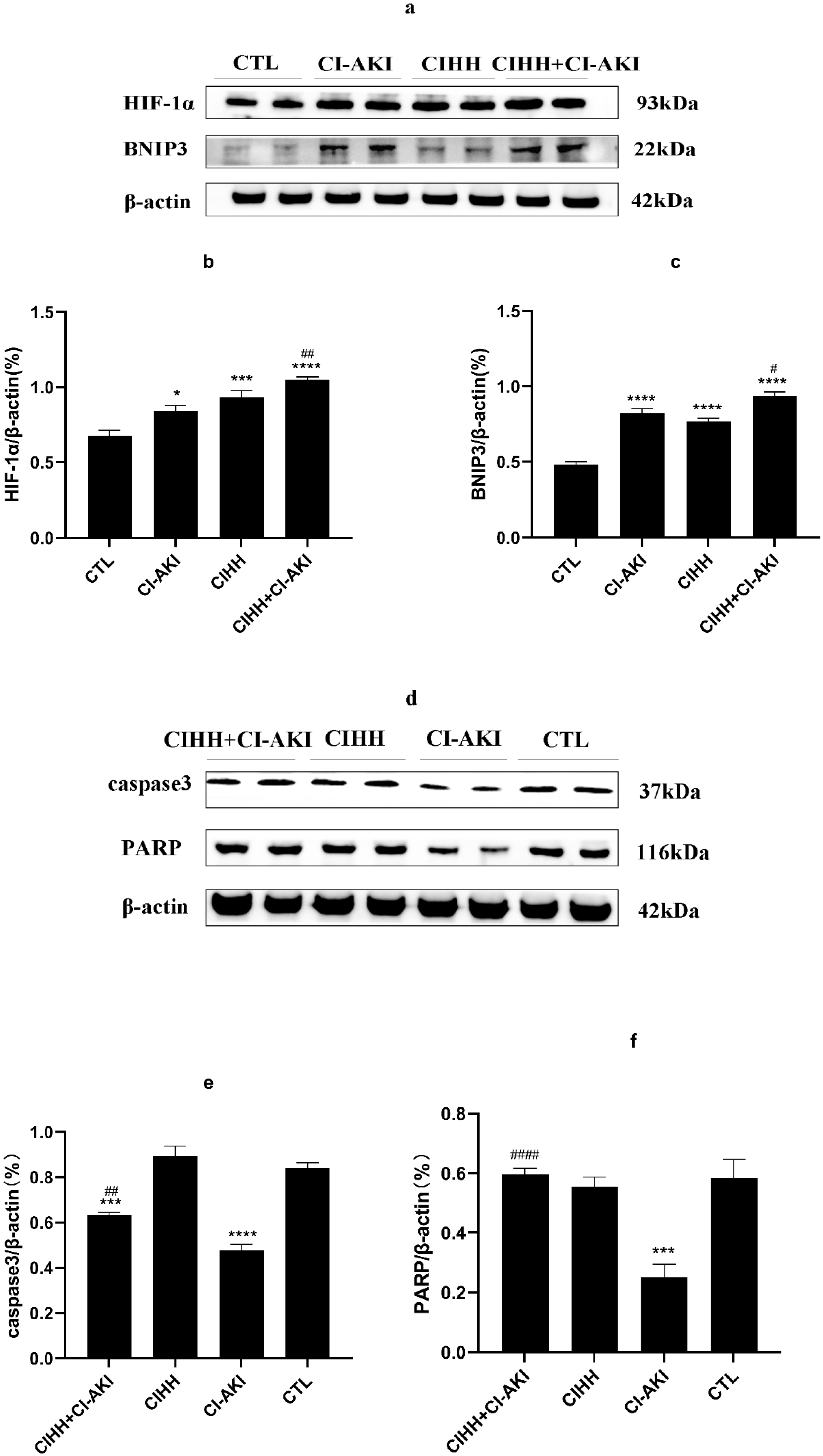
Effects of CIHH on HIF-1α,BNIP3,caspase3 and PARP proteins expression in renal of CI-AKI. afrom the same gel. HIF-1α,BNIP3,β-actin in the picture a from the same gel. Caspase3,PARP,β-actin in the picture d from the same gel Representative blots of HIF-1α,BNIP3,caspase3 and PARP proteins expression(a,d). Analyzed data of HIF-1α ,BNIP3 ,caspase3 and PARP proteins expression(b,c,e,f). CTL, normal-controlled group; CI-AKI, contrast induced acute kidney injury group; CIHH, chronic intermittent hypobaric hypoxia pretreatment group; CIHH+CI-AKI, contrast induced acute kidney injury group with chronic intermittent hypobaric hypoxia pretreatment. Dates were presented as means ± SEM (n=6).*P<0.05,***P<0.001,****P<0.0001versus.CTL;#p<0.05,##p<0.01,####p<0.0001versus.CI-AKI. The uncropped blots are presented in Supplementary Figure.

## 3. Discussion

The incidence of CI-AKI has decreased in recent years with the use of isotonic/hypotonic contrast agents, but the number of CI-AKI cases is still increasing. In the general population, the incidence of CI-AKI is 2.0%, but in the population with comorbid chronic kidney disease, diabetes, advanced age, and emergency percutaneous coronary intervention (PCI), the incidence of CI-AKI rises to 30%[24]. Although studies have shown that some drugs or methods have a preventive effect on CI-AKI, such as lignans[25], acetylcysteine[26], sodium bicarbonate[27], hydration[28]etc. But, there is still no consensus on these methods. Once CI-AKI occurs, there is no effective treatment. Therefore, it is particularly important to prevent the occurrence of CI-AKI.CIHH has been shown to have many benefits to the organism both in the exercise business and in animal experiments. The present study is the first to suggest that CIHH pretreatment before administration of contrast media can prevent or attenuate the occurrence of CI-AKI in rats.

In the present study, the rats in the CI-AKI group showed increased heart rate, elevated mean arterial pressure, significantly increased blood creatinine and blood urea nitrogen, aqueous degeneration of renal tubular epithelial cells, eosinophilic masses in the lumen of a large number of renal tubules, tubular necrosis, and unclear tubular lumen structure, suggesting that the rats in the contrast nephropathy model had significant renal injury. Although the rats blood creatinine and blood urea nitrogen levels in the CIHH prevented CI-AKI group were still higher than the baseline values, their elevation was significantly reduced, and blood pressure and heart rate were significantly lower than those in the CI-AKI group, while the renal pathological injury was significantly reduced. These suggest that CIHH has a protective effect against contrast-induced acute kidney injury.

ROS plays an important role in the development of contrast nephropathy, which can cause direct damage to vascular endothelial cells and renal tubules, and in turn aggravates the hypoxic damage to renal parenchyma caused by vascular endothelial dysfunction and dysregulation of renal tubular transport[6]. SOD is an enzyme that can restore the balance of cellular redox disorders caused by local ischemia and hypoxia, and SOD catalyzes the disproportionation of O_2_ radicals in H_2_O_2_, which can prevent the accumulation of reactive oxygen species and play a role in scavenging oxygen free radicals[29]. In the kidney tissue of contrast nephropathy, ROS is increased, SOD activity is decreased, and MDA levels are increased[23, 30]. Several studies have shown that increased SOD activity attenuates tissue or cellular damage caused by oxidative stress[31–34]. In the present study, renal SOD activity was significantly lower and MDA levels were significantly higher in the CI-AKI group, which is consistent with other related studies. Renal SOD activity was significantly higher and MDA levels were lower in the CIHH prevented CI-AKI group than in the CI-AKI group, indicating that CIHH pretreatment could protect kidney by attenuating renal oxidative stress.

Although the pathogenesis of CI-AKI remains unclear, ischemia and hypoxia play an important role in the development and progression of contrast nephropathy[2]. More than 100 genes have been identified to cause a significant increase in HIF-1α in response to renal hypoxia[35]. By upregulating of renal HIF-1α levels could ameliorate renal fibrosis and glomerular mesangial hyperplasia in diabetic rats[10]. Recent studies have shown that activation of the HIF-1α/BNIP3 signaling pathway under hypoxic conditions results in enhanced mitochondrial autophagy, which can have a protective effect on tissues or cells. Lin et al. showed that renal injury and apoptosis were exacerbated in BNIP3 knockout mice given contrast agents and in HK2 cells transfected in advance with si-BNIP3 to silence BNIP3[8].In a study by Fu et al, inhibition of mitochondrial autophagy in the kidneys of HIF-1α knockout ischemia/reperfusion mice exacerbated tubular apoptosis and renal injury, and increased BNIP3 expression in the kidneys of knockout mice resulted in enhanced mitochondrial autophagy, reduced tubular apoptosis and renal injury, indicating that the HIF-1α/BNIP3 signaling pathway mediates ischemia/reperfusion injury and that activation of this pathway may protect against ischemia/reperfusion injury[7].In the present study, the renal HIF-1α and BNIP3 protein expression levels were increased in the CI-AKI group compared to CTL group, and CIHH prevented CI-AKI group were significantly higher compared to CI-AKI group, indicating that CIHH preadaptation could enhance mitochondrial autophagy through activation of the HIF-1α/BNIP3 signaling pathway to protect against contrast-induced renal injury.

Not only the toxic effects of the contrast agent itself, but also ischemia and hypoxia, oxidative stress injury and apoptosis play an important role in the development and progression of CI-AKI. Several studies have shown that caspase3 activity increases and the level of anti-apoptotic factor BCL-2 decreases in acute kidney injury, leading to increased apoptosis, which caused renal tubular epithelial cell injury, and that administration of drugs to inhibit aspase3 activity can reduce cell apoptosis and thus improve kidney injury[25, 36–38].Caspase3 was inactive, but its activated product, cleaved-caspase3,which could activate PARP caused apoptosis. Romano et al. showed that caspase3 and PARP levels were reduced and apoptosis was increased in a mouse model administered with contrast agents suggesting that caspase3-dependent apoptotic pathway mediates iodine contrast-induced acute kidney injury[39].In the present experiment, caspase3 and PARP levels in the kidney tissues of rats in the CI-AKI group were significantly lower compared to CTL group, but the levels of caspase 3 and PARP in the CIHH prevented CI-AKI group were significantly increased compared to CI-AKI group, indicating that CIHH pretreatment played a protective role against contrast-induced acute kidney injury by increasing the anti-apoptotic capacity of renal cells.

## 4. Materials and methods

### 4.1 Animals and CIHH pretreatment

Male Sprague-Dawley (SD) rats, SPF grade, 24 in total (weight 160±10 g, buy from Hebei Ivivo Biotechnology Co., Ltd.), were purchased and kept in the laboratory of Hebei Medical University, where the laboratory temperature was constant at 22±1°C and received light/dark time for 12 h each, with free access to water and food. At the beginning of the experiment, 24 male SD rats were randomly divided into 4 groups of 6 rats each: normal-controlled group (CTL, rats were kept in the laboratory environment at all times), CI-AKI group (rats were kept in the laboratory environment at all times), CIHH group (SD rats were placed in a hypobaric hypoxic chamber simulating 4000m altitude for 5h per day for 35 days), CIHH prevented CI-AKI group ( SD rats were placed in a hypobaric hypoxic chamber simulating 4000m altitude for 5 hours per day for 35 days before CI-AKI modeling). The experiment was approved by the ethics committee of Hebei Provincial People’s Hospital and completed by the Department of physiology of Hebei Medical University. The study was carried out in compliance with the ARRIVE guidelines.

### 4.2 CI-AKI rats model establishment

The modeling method was referred to Bae et al.[22], as follows: rats in the CI-AKI group and CIHH to prevent CI-AKI group were sequentially injected with indomethacin (10mg/kg, Aladdin,China,I106885-5g, purity98%), Nω-nitro-L-arginine methyl ester (10mg/kg, 15 min later, Sigma, USA, N5751-5G, purity>98%), and ioversol injection (8.3 mL/kg, 30 min later , 350mgI/ml, Jiangsu Hengrui Pharmaceutical Co., Ltd, China) through the tail vein, and rats in the CTL and CIHH groups were injected with saline through the tail vein 0.5mL/time,3 times in total. After injection, all rats were kept in the laboratory environment and resumed free access to water and food.

## 5. Detection methods of various indicators

### 5.1 Heart rate and mean arterial pressure measurements

Heart rate and mean arterial pressure were measured by a tail cuff manometer (LE5001, Panlab) every morning (7:00-10:00 am) from one week before injecting drug until 24 hours after contrast agent injection, or before blood collection if blood specimens were to be collected from the tail vein.

### 5.2 Testing of biochemical indicators

Blood was collected from the tail vein of rats before and 24 hours after contrast agent injection, respectively. Then centrifuged them for 15 min at 4°C and 3000 rpm using a tabletop high-speed refrigerated centrifuge (DaLong, D3024R), and the supernatant was taken for the determination of serum creatinine and serum urea nitrogen by an automatic biochemical analyzer (Shenzhen Redu Life Science, Chemray800).

### 5.3 Measurement of Biomarkers of Oxidative Stress

The rats were killed by neck leading at 24 h after contrast agent injection, and the isolated right kidney tissues were washed with saline (pre-cooled on ice in advance) to remove residual blood and homogenized, then using a benchtop high speed frozen centrifuge (DaLong, D3024R) to centrifuge for 15 min at 4°C, 3000 rpm /min,, and the supernatant was collected. SOD activity and MDA levels were determined by using two commercial experimental kits (Nanjing Jiancheng Institute of Biological Engineering) according to the assay instructions of the kits[23].

### 5.4 Histological examination

Rats were executed 24 hours after drug injection, and the left kidney was fixed in 4% paraformaldehyde solution for 48 hours and then dehydrated in gradient alcohol, transparent in xylene, and embedded in paraffin wax. The paraffin kidney tissue block was cut into thin slices of approximately 3μm thickness using a microtome (Leica, Germany), then stained with hematoxylin eosin staining (HE staining), periodic acid Schiff reaction (PAS)and Masson trichrome after dewaxing in xylene and rehydrating in gradient alcohol. Last observed and photographed in positive White light microscope (Nikon (Japan), Eclipse Ci-L), and each tissue section was carefully observed under the microscope and histological changes were recorded.

### 5.5 Immunoblot analysis

Rats were executed 24 h after drug injection, the right kidney was taken and cut with tissue scissors. The kidney tissues were homogenized in tissue/cell lysate. Protein samples were separated by SDS–PAGE, transferred to a PVDF membrane (Millipore Corporation, USA) that was blocked for 1h with 5%(w/v) non-fat milk in Tris-buffered saline, and incubated with antibodies against HIF-1α (1:500, Wanleibio, China, WL01607), BNIP3 (1:1500, Wanleibio, China, WL01139), caspase3 (1:1500, Affinity, USA, AF6311)), PARP (1:750, Wanleibio, China, WL01932), overnight at 4 °C on a constant temperature refrigerator shaker. The same membrane was stripped and re-blotted with an β-actin antibody (1:5000, Proteintech, USA, 20536-1-AP) for normalization. Blots were developed by the chemiluminescent detection method (Amer sham ECL). The protein blots were quantified by densitometry using Image J software and normalized to β-actin.

### 5.6 Data analysis

All data were performed with GraphPad Prism 9.0.0,121,software. All groups with n=6 each were analyzed by one-way analysis of variance (ANOVA). Results are presented as mean±SEM and P<0.05was considered significant.

## Data availability statement

The data sets generated and / or analyzed during this study are not public due to the data store in Baidu online disk, but Baidu online disk is not a public network in the world, not everyone can relevant information links can be obtained, but can be obtained from the corresponding author upon reasonable request.

## Original data owner declaration

Each person who needs this data can obtain relevant data from the author Kai-min Yin and contact the.

## Acknowledgements

We thank Prof. Hui-jie Ma and his group in the Physiology Department of Hebei Medical University for providing the Hypobaric hypoxic chamber for this study and for their help in the experiments.

## Author Contributions

Kai-min Yin made a major contribution to this paper.

